# Origin, domestication, and diversity of the climate resilient Ethiopian cereal teff

**DOI:** 10.1101/2025.09.07.674720

**Authors:** CM Wai, ML Wilson, E Braun, J Baczkowski, S Chanyalew, M Dell’Acqua, E Grotewold, A Thompson, R VanBuren

**Affiliations:** Plant Resilience Institute, Michigan State University, East Lansing, MI 48824, USA; Department of Horticulture, Michigan State University, East Lansing, MI 48824, USA; Department of Plant Biology, Michigan State University, East Lansing, MI 48824, USA; Department of Biochemistry and Molecular Biology, Michigan State University, East Lansing, MI 48824, USA; Department of Plant, Soil, and Microbiology, Michigan State University, East Lansing, MI 48824, USA; Ethiopian Institute of Agricultural Research, PO Box 2003, Addis Ababa, Ethiopia; Institute of Plant Sciences, Scuola Superiore Sant’Anna, Pisa, Italy

**Author notes:** Corresponding author: Robert VanBuren. These authors contributed equally to this work (CM Wai, ML Wilson).

## Abstract

Teff (*Eragrostis tef*) is a climate-resilient cereal and staple crop in the Horn of Africa, cultivated by millions of smallholder farmers. Despite its cultural and nutritional significance, teff has limited genomic tools, constraining progress in yield improvement and trait optimization. Here, we assembled and resequenced the Teff Association Panel (TAP), a diverse set of 265 landraces and cultivars, to investigate the domestication history, genetic diversity, and agronomic traits of teff. Phylogenetic and population genetic analyses confirm *Eragrostis pilosa* as the direct progenitor of teff, with domestication likely occurring in the Tigray region of Ethiopia. We generated a high-quality reference genome for *E. pilosa* and found extensive chromosomal collinearity and minimal gene loss compared to teff, with teff-specific genes enriched in functions related to domestication traits. We identified signatures of selection across the teff genome and used genome-wide association studies (GWAS) to map loci associated with seed color and panicle architecture. Integrated GWAS, expression, metabolic, and comparative genomic analyses pinpoint a major seed color locus containing orthologs of the *TT7* cytochrome P450 gene involved in flavonoid hydroxylation and the *TT2* MYB transcription factor regulating proanthocyanidin biosynthesis. For panicle architecture, we identified a strong association near a teff ortholog of Dense and Erect Panicle 1 (DEP1), a key regulator of inflorescence structure in rice and sorghum. This suggests convergent selection on shared genetic pathways controlling panicle morphology across cereals. Our findings confirm the evolutionary origin of teff, identify key domestication loci, and highlight untapped genetic diversity among Ethiopian breeding lines. These genomic resources provide a foundation for accelerating teff improvement through molecular breeding and genome editing to enhance yield, resilience, and nutrition.

## Introduction

Humans domesticated dozens of cereals within the Poaceae, or grass family, over the past 12,000 years, and cereals are a cornerstone of global food security. Cereals were independently domesticated across each of the historical centers of crop origin from phylogenetically diverse and ecologically adaptable wild grasses. Despite their diverse origins, the domestication of cereals converged on a common set of traits for enhanced yield, harvestability, and nutrition, using a remarkably conserved genetic architecture ^1^. This includes key genetic loci for reduced seed shattering, increased grain size, and changes in whole plant and panicle architecture. However, crop domestication is a long and complex process ^2^, and early cereals were shaped by repeated introgressions from wild populations, which contributed genetic variation that supported selection for locally adapted traits governed by more complex genetic architectures ^3^.

Millets, a subset of cereals, are characterized by their small but nutritious grains and remarkable adaptability to arid or semi-arid regions ^4^. Their domestication history is rooted in indigenous agricultural practices, and they are primarily cultivated in diverse, small-scale cropping systems that sustain ∼250 million farmers worldwide. Unlike major cereals, classic domestication traits, such as non-shattering seeds, lodging tolerance, and increased grain size are often incompletely fixed in millets, contributing to lower yield under optimized conditions. Despite these challenges, millets have undergone intensive selection for local adaptations that promote stable and consistent yields under marginal growing conditions. This resilience makes millets vital for developing sustainable, diversified, and climate-resilient agricultural systems, although significant challenges remain in optimizing their production and addressing yield gaps.

Teff (*Eragrostis tef*), a small-grained millet native to the Horn of Africa, is a cornerstone of Ethiopian agriculture, where it is grown on ∼24% of the total cultivated land and contributes to 21% of the yearly grain production ^5,6^. Teff provides an estimated two-thirds of the daily protein intake for most Ethiopians and it is a significant economic commodity, as its market price is often two to three times higher than maize ^5,7^. Cultivated by 6.2 million small-scale farmers, teff exhibits extensive local adaptation, with an estimated 5,000 landraces distributed across Ethiopia’s diverse growing regions ^8,9^. Teff adaptation to this broad environmental variation enables teff production across different agro-ecological zones, including regions where major cereals may fail.

Ethiopia produces ∼90-95% of the world’s annual 4-5 million metric tons of teff, but recently the cereal has grown in popularity elsewhere in the world because of its superior nutritional profile, gluten free grain, and climate resilience ^10^. Despite its importance, research and development for teff have historically been underfunded, limiting global awareness and access to the crop ^10^. As a primarily self-pollinating grass with outcrossing rates as low as 1–2% ^11^, breeding and introgressing traits is a major challenge in teff. However, CRISPR/Cas9 editing of the SD-1 gene in teff has successfully conferred semidwarfism by reducing culm and internode lengths, significantly improving lodging resistance without affecting panicle or peduncle lengths ^12^. This gene editing platform, along with advanced genomic resources ^13–15^, provide a foundation for the rapid improvement of teff, but critical gaps remain in our understanding of the genetic diversity, domestication history, and the genetic basis of key agronomic traits.

Here, we explore the origin, genome evolution, and domestication of teff. Using a newly constructed Teff Association Panel (TAP), we investigate the genetic and evolutionary history of teff domestication and improvement. Sequencing of wild Eragrostis germplasm confirms the direct progenitor of teff, and through comparative genomics, we identify changes in genome architecture associated with domestication. We use the panel to identify genetic loci associated with seed color and panicle architecture, and identify a co-selected locus associated with domestication. These findings, combined with advanced genomic resources, provide a robust foundation for accelerating teff breeding to address critical agronomic challenges and meet future demands.

## Results

### Cataloging the genetic diversity of teff

To explore the genetic diversity of teff, we assembled a panel of 387 landraces, breeding lines, elite cultivars, and wild *Eragrostis* germplasm from the USDA Germplasm Resources Information Network (Supplemental Table 1). Accessions were resequenced and aligned to the ‘Dabbi’ reference genome ^13^, with median read mapping rates of 98.8% for teff accessions, 95.0% for the putative wild progenitor *Eragrostis pilosa*, and 41.3% for all other Eragrostis species (Supplemental Table 1). Across the panel, we identified ∼8.1 million single nucleotide polymorphisms (SNPs), and 1.5 million insertions/deletions (indels). The USDA GRIN germplasm was collected in the 1950s-1980s during the establishment of the Plant Genetic Resources Center of Ethiopia, and many accessions are duplicated because of insufficient passport data on origin and local variety names ^16^. Using identity-by-descent, we found that 135 of the 363 sequenced teff lines were cryptically related and correspond to duplicated Ethiopian varieties. We retained the accession with the highest sequence coverage, resulting in a final panel of 265 unique teff lines. We refer to this refined set of germplasm as the Teff Association Panel (TAP), and we used the sequence variation within these lines and the 24 wild Eragrostis accessions for downstream analyses.

Teff is cultivated by millions of small-scale farmers in Ethiopia, and there is tremendous diversity found across the thousands of locally adapted and farmer-selected varieties. The Ethiopian Biodiversity Institute maintains the largest collection of teff, with approximately 6,000 accessions that encompass the global diversity of local landraces, cultivars, and lines used for teff breeding ^17^. To assess how much of this diversity the USDA germplasm captures, we compared the genetic variation of our resequencing data to the Ethiopian Teff Diversity Panel (EtDP), 321 farmer varieties sourced from the Ethiopian Biodiversity Institute. The EtDP spans the geographical and agroecological range of teff and reflects the genetic diversity preserved within Ethiopia ^14^. Using a common set of 7,747 SNP based markers present in both panels, we identified nine distinct subpopulations using ADMIXTURE, with each group having similar representation in the TAP and EtDP (Figure 1e)^18,19^. These subpopulations comprise 37 to 105 teff accessions each and have high levels of admixture that reflect limited genetic stratification. Principal component analysis separates the samples by subpopulation, and accessions from both the EtDP and TAP are widely distributed across the PC1 and PC2 space, confirming a comprehensive representation of global teff diversity in both panels (Figure 1a).

**Figure 1.**
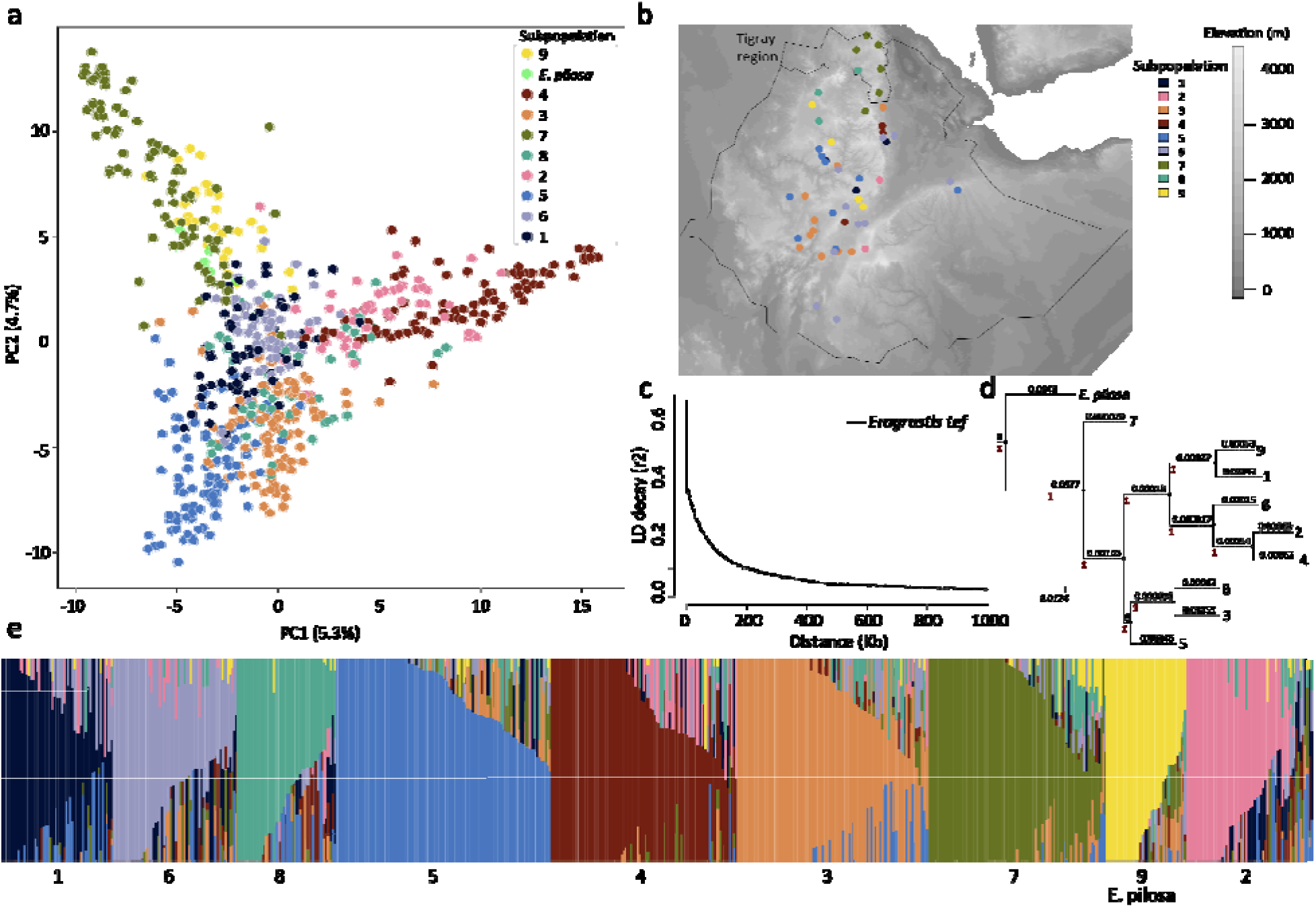
Genetic diversity of teff and its wild progenitor *Eragrostis pilosa*. (a) Principal component analysis of commonly genotyped SNPs between the EtDP and TAP lines. Accessions are colored by the nine subpopulations identified in the ADMIXTURE analysis. The *E. pilosa* accessions are found in subpopulation 9, but are colored separately here. (b) Distribution of georeferenced lines from the TAP across an altitudinal map of Ethiopia. The putative center of origin for teff, the Tigray region in Northern Ethiopia is outlined. (c) Linkage disequilibrium decay plot teff lines in the TAP using the full set of genome wide variants. (d) Phylogenetic network inferred by Treemix of the relationships between the teff subpopulations and *E. pilosa* with *E. heteromera* as an outgroup. The drift parameters for each branch are shown. (e) ADMIXTURE results for the common SNPs between the EtDP and TAP. Each accession is represented by a vertical line, and colored by the proportion of each subpopulation in the genome of each line.

Accessions from the EtDP and TAP originate from all the major teff-growing regions in Ethiopia, and we observed some degree of geographic separation among subpopulation (Figure 1b). Despite this, most subpopulations exhibit broad and overlapping geographic distributions, underscoring the extensive admixture and minimal genetic divergence within teff. This is further supported by the low fixation index (Fst), a measure of genetic variation between populations, averaging 0.07 across all pairwise comparisons of the nine subpopulations, indicating slight genetic separation between teff subpopulations (Supplemental Figure 1). We confirmed a geographically distinct subpopulation (subpopulation 7) from the Tigray region of Northern Ethiopia, the hypothesized center of origin for teff ^20,21^. This subpopulation has a higher pairwise Fst than the other subpopulations, suggesting a unique genetic composition (Supplemental Figure 1). This distinct germplasm from Tigray may have been shaped by early domestication events and recurrent genetic exchange with wild progenitors. Breeding lines curated by the Ethiopian Biodiversity Institute are predominantly found within subpopulations 2 and 8, indicating that a vast reservoir of genetic diversity is underutilized in teff breeding programs. This suggests an opportunity to broaden the genetic base of teff cultivars, potentially enhancing traits such as yield, resilience, and nutritional content ^14^. This strategic expansion of the genetic pool could be pivotal for future teff improvement efforts, catering to both national and global demands for this important crop.

Teff was domesticated in the Northern Ethiopian Highlands, likely from the wild grass *Eragrostis pilosa* ^20,22,23^. While teff and *E. pilosa* are capable of producing fertile interspecific hybrids and share similarities in numerous traits, prior studies have had too few markers or polymorphic sites to verify the origin of teff. We compared whole genome data for 12 *E. pilosa*, and other hypothesized teff progenitors including *E. heteromera*, *E. macilenta*, *E. mexicana*, and *E. papposa* to teff. Genetic analyses including admixture, phylogenetic inference, dimensionality reduction, and population differentiation (Fst) provide substantial support that *E. pilosa* is the direct ancestor of teff (Figure 1, Supplemental Figure 1). Maximum likelihood phylogenetic analysis using TreeMix, with *E. heteromera* as an outgroup, positions *E. pilosa* as sister to teff and highlights an early divergence of accessions from Tigray within the teff lineage (Figure 1d). *E. pilosa* samples do not form a distinct group in the ADMIXTURE analysis but are instead integrated within the genetic makeup of teff, primarily clustering within subpopulation 9, with notable admixture observed from subpopulation 7 (Figure 1e). This integration is further supported by the principal component analysis, which clusters *E. pilosa* alongside teff germplasm from both the Tigray region (subpopulation 7) and subpopulation 9, suggesting a genetic continuity between these groups (Figure 1a). The average pairwise Fst between *E. pilosa* and teff subpopulations is 0.3, which is consistent with expected genetic divergence between a crop and its direct progenitor (Supplemental Figure 1). The lowest Fst values were observed between *E. pilosa* and subpopulations 7 and 9. Collectively, these analyses highlight the close genetic relationship between teff and *E. pilosa*, strongly supporting the hypothesis that *E. pilosa* served as the primary ancestor of teff.

### Exploring the evolutionary origin of teff

To gain a deeper insight into the origin and domestication history of teff, we constructed a d*e novo* reference genome of the putative progenitor *E. pilosa* (Figure 2a), and explored genome evolution between these grasses. We generated 64.5 gigabases of High Fidelity PacBio (HiFi) long-read sequencing data and assembled the reads using Hifiasm ^24^. The resulting *E. pilosa* genome assembly has 70 contigs with a total length of 560 Mb and an N50 of 14.7 Mb. Most *E. pilosa* chromosomes are assembled into 2-3 contigs and the total assembly size is similar, but ∼17 Mb smaller than the ‘Dabbi’ teff reference ^13^. Using the MAKER pipeline for *ab initio* gene prediction, we identified 69,668 gene models in *E. pilosa*, which is comparable to the 68,255 gene models annotated in ‘Dabbi’.

**Figure 2.**
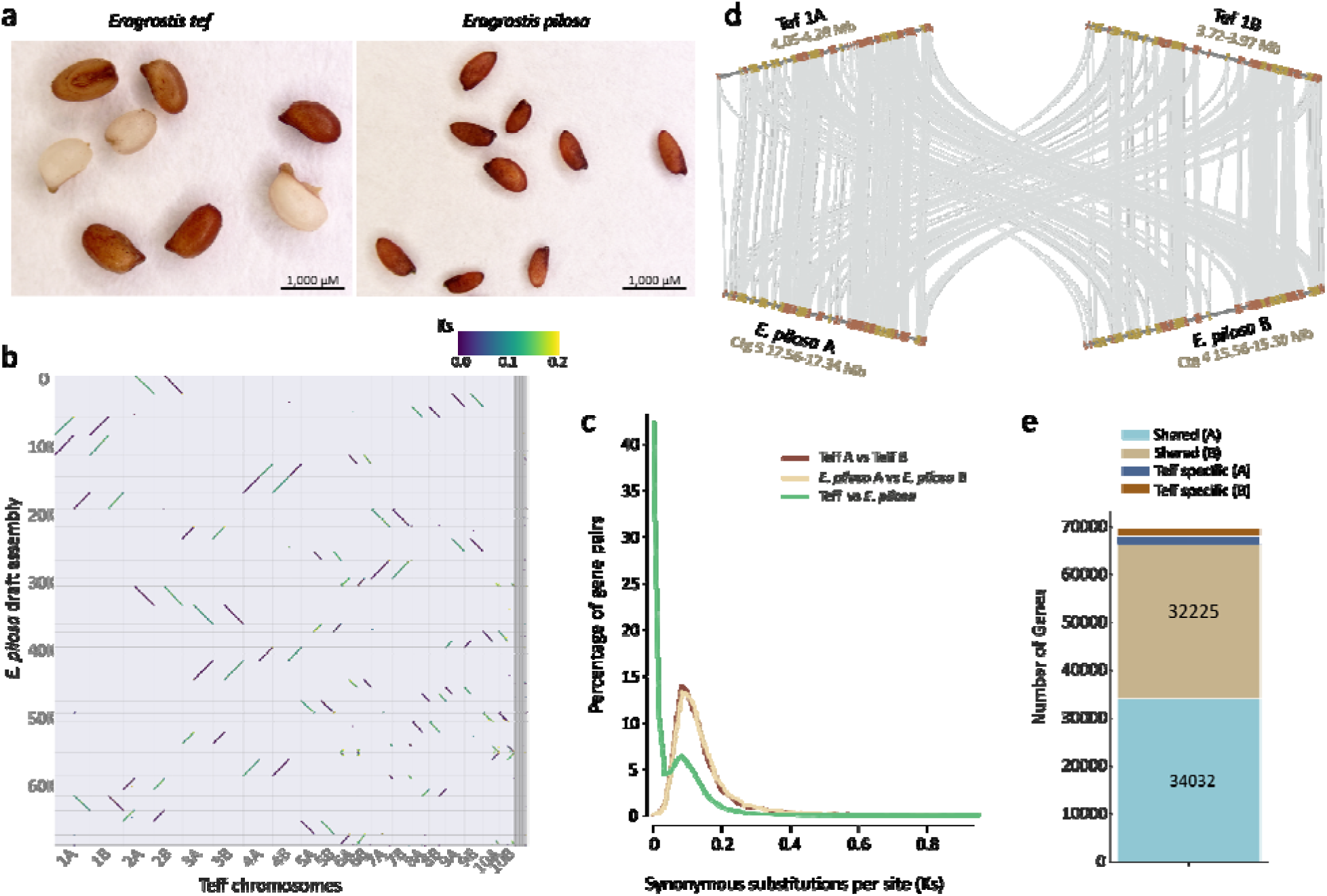
Comparative genomics of the teff and *E. pilosa* genomes. (a) Comparison of mixed brown and white teff seeds (left) and brown *E. pilosa* seeds (right). (b) Macrosyntenic dot plot between the *E. pilosa* and teff genomes where each dot represents a syntenic gene pair and dots are colored by the Ks of each gene pair. (c) Histogram of Ks for syntenic gene pairs between *E. pilosa* and teff (green), homeologs between the teff A and B subgenomes (purple) or *E. pilosa* A and B subgenomes (blue). (d) Microsynteny between the teff and *E. pilosa* genomes. A portion of Chromosomes 1A and B are shown where individual genes are shown in orange or gold and syntenic gene pairs are connected by gray lines. (e) Stacked bar plot showing the gene pairs conserved between teff and *E. pilosa* and genes unique to teff for the A and B subgenomes.

The A and B subgenomes of teff exhibit clear orthology to two subgenomes of *E. pilosa*, with broadly conserved macrosynteny, and no structural rearrangements (Figure 2b). To characterize the *E. pilosa* polyploidy event, we calculated synonymous substitution rates (Ks) between homoeologous gene pairs within *E. pilosa* and syntenic orthologs across the two species. Homoeologs within the A and B subgenomes of *E. pilosa* have a median Ks of 0.14, similar to the Ks of teff A and B homoeologs (0.15), supporting that the allotetraploidy event predates teff domestication and is shared with *E. pilosa* (Figure 2b, c). Consistent with this, the median Ks values for syntenic gene pairs between the teff A and *E. pilosa* A subgenomes and the teff B and *E. pilosa* B subgenomes are 0.0036 and 0.0039, respectively (Figure 2c). Based on Ks, we assigned 21 and 22 contigs to the A and B subgenomes of *E. pilosa*, respectively (Figure 2b). Using a widely accepted mutation rate for grasses (1.5 × 10^−8 substitutions per nonsynonymous site per year), we estimate that this accession of *E. pilosa* and teff diverged approximately 120,000 years ago, and confirm the Eragrostis allotetraploidy event occurred ∼5 million years ago ^13^.

Teff and *E. pilosa* have compact genome architectures, with a monoploid genome size of 280-300 Mb and a relatively low proportion of repetitive elements. Repeats span 33% of the *E. pilosa* genome, with gypsy long terminal repeat retrotransposons (10.9%) and Heilitron transposons (8.5%) being the most abundant elements (Supplemental Table 2). Repeat element composition is similar between teff and *E. pilosa*, but helitrons are notably more abundant in *E. pilosa* compared to teff (8.5% vs 2.3%). Helitrons preferentially integrate into genic regions, influencing gene expression and function ^25^, and the lower abundance of helitrons in teff could reflect selection for genomic stability during domestication.

The teff and *E. pilosa* genomes have conserved gene content and order across the A and B subgenomes (Figure 2d). Roughly 95% of teff genes have syntenic orthologs in *E. pilosa,* with 1,702 and 1,751 teff genes having no orthologs in the corresponding *E. pilosa* A and B subgenomes, respectively (Figure 2e). There is no bias fractionation or significant difference in gene loss between A and B, consistent with previous observations of subgenome stability in teff^13^. The proportion of conserved genes between teff and its wild progenitor is considerably higher than other wild and domesticated cereals such as maize, sorghum, and rice, where half or more genes are dispensable ^26–28^. This unusual conservation could be explained by the compact genomes, low transposable element content, high selfing rate, and broad genome stability observed across sequenced chloridoid grasses ^29–32^. The 3,453 teff genes that are absent from *E. pilosa* are enriched in functional roles related to core and specialized metabolism, stress responses, and metal ion transport (Supplemental Figure 2), and could be linked to selection during domestication.

### Signatures of teff domestication and improvement

Despite the vast diversity of morphological and agronomic traits, previous marker-based studies have identified limited genetic variation across teff germplasm ^33^. Using invariant sites of reads mapped the ‘Dabbi’ reference genome, we calculated the nucleotide diversity (π) within the TAP and *E. pilosa*. The genome-wide nucleotide diversity of all teff accessions is 9.7 x 10−4 which represents a threefold reduction in diversity compared to *E. pilosa* (π = 2.9 x 10-3). Nucleotide diversity within subpopulations of teff is similar (mean = 8.9 x 10−4), but the subpopulation 7 from Tigray has the highest π of 1.1 x 10-3. Teff has higher nucleotide diversity than Fonio (π = 6.2 × 10–4) ^34^ but is less diverse than other millets including foxtail millet (π = 1.6 × 10–3) ^35^, pearl millet (π = 2.3 × 10–3)^36^, and sorghum (π = 2.4 × 10–3) ^37^. Nucleotide diversity is ∼15% higher in the A subgenome compared to B in both teff and *E. pilosa*, which is a similar pattern to other allopolyploids including wheat ^38^, and barnyard millet ^39^, and likely represents differences in evolutionary history and selection pressures of the subgenomes.

To identify potential selective sweeps during teff domestication, we looked for signatures of reduced nucleotide diversity (π*_E. pilosa_* /π_teff_), elevated cross-population differentiation (XP-CLR)^40^, and local distortions in the site frequency spectrum using RAiSD (μ statistic) ^41^ in teff and *E. pilosa* (Figure 3; Supplemental Figure 3). We calculated values in 25kb sliding windows across the teff genome and identified putative selective sweeps based on overlapping regions found in the top 5% of each statistic. We identified 78 putative selective sweeps spanning 5.2 Mb or 0.9% of the teff genome, with an average size of 67Kb (Figure 3). Sweeps are distributed unevenly across the subgenomes, with 50 regions spanning 3.4 Mb in the A and 28 regions spanning 1.8 Mb of the B subgenome. Putative swept regions contain 731 genes (Supplemental table 3). These genes span a broad functional space but show notable enrichments in stress response and hormone signaling, including NB-ARC domain-containing disease resistance genes, heat shock proteins, and components of auxin and cytokinin pathways. Several transcription factor families (e.g., WRKY, NAC, GRAS, and AP2/ERF) are also within swept regions, and could be related to regulatory shifts in growth, development, and stress responses. Genes involved in cell wall biosynthesis, sugar transport, and oxidative stress tolerance were also common, suggesting possible selection for traits related to architecture, lodging resistance, and resilience.

**Figure 3:**
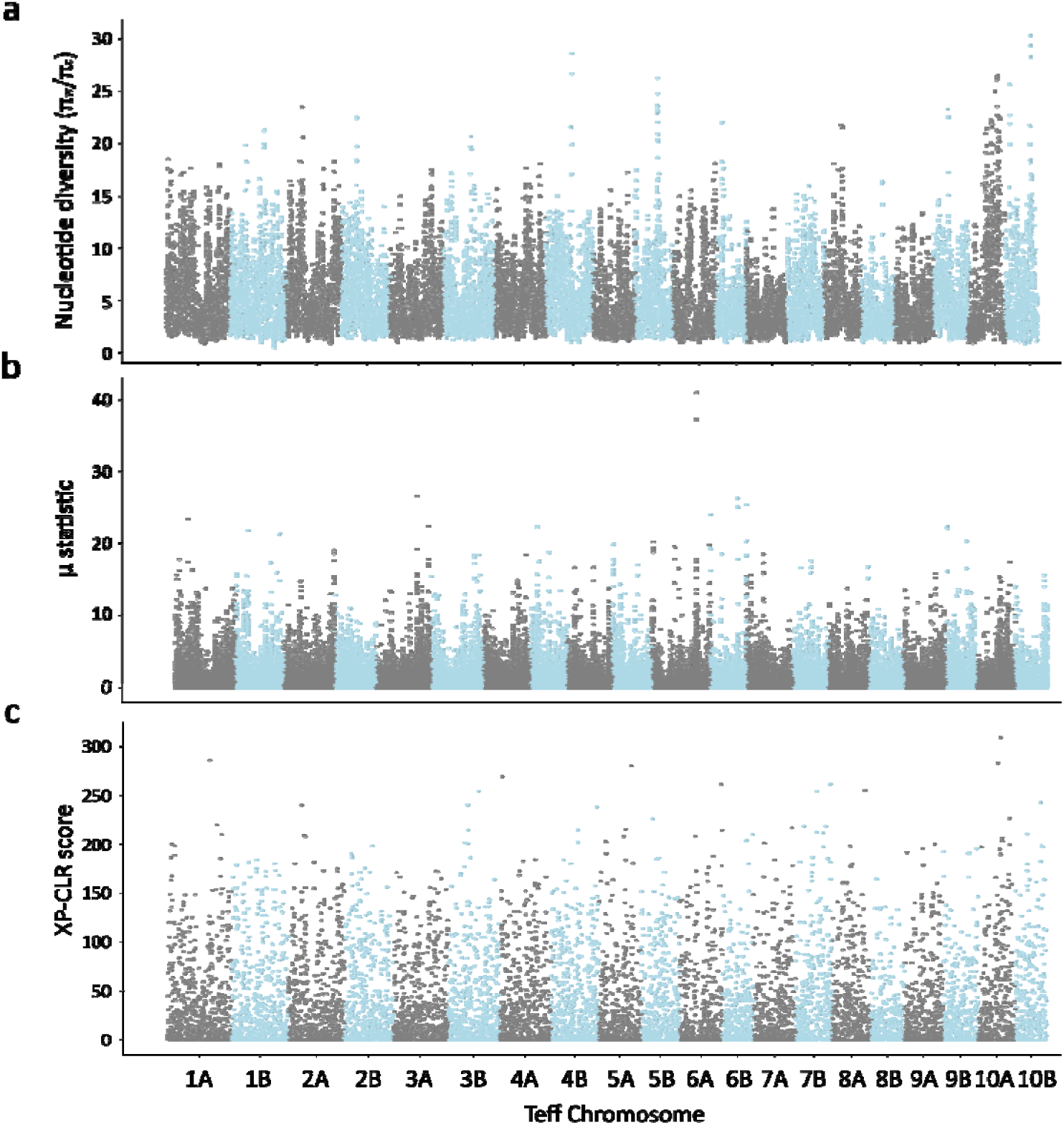
Detecting Selective sweeps in the teff genome. Genome-wide scan for selective sweeps in teff using three complementary approaches: (a) π ratio (π *E. pilosa* / π teff), (b) XP-CLR, and (c) RAiSD μ-statistic. Analyses were conducted in 25 kb sliding windows across the 20 teff chromosomes. Higher values in each panel indicate putative regions under selection during domestication or improvement. Shaded bars alternate between chromosomes for visual clarity.

### Agronomic and domestication traits in teff

Teff has a rich and complex recent evolutionary history shaped by the diverse trait preferences of small-scale farmers in Ethiopia and the varying climatic and agroecological conditions they encounter. Farmer preferences depend on the end use of the grains or fodder, as well as factors such as geography, climate, and gender-specific roles in agriculture ^14^. As a result, teff varieties display remarkable variation in plant and panicle architecture, seed color and size, maturity time, and yield-related traits. To investigate the diversity and genetic basis of domestication traits in teff, we phenotyped the TAP over successive field seasons in Michigan, and performed genome-wide association studies (GWAS) on key cereal traits.

Agronomic and nutritional traits are highly diverse within the TAP. Panicle architecture, lodging tolerance, and seed color show substantial variation both within and across teff subpopulations, with only a few differences in trait distribution clearly delineating groups. Compared to its wild progenitor, *E. pilosa*, teff displays clear domestication signatures including larger seeds with varying colors, taller plants with fewer tillers, more closed panicles, and reduced seed shattering. Most of the improved teff varieties from the Ethiopian Institute of Agricultural Research breeding program are in subpopulation 2, which has exclusively white seeds (Supplemental Table 4), while other subpopulations exhibit a mix of white, brown, or mixed seed colors. White-seeded teff is generally preferred by consumers ^42,43^ and is more widely grown in Ethiopia due to its higher market value. Brown teff is traditionally consumed by subsistence farmers, but has seen increased demand in Ethiopia and globally for its high iron content and superior nutritional profile ^44^. Brown-seeded varieties sometimes have greater yield at high altitudes and are generally viewed as more resilient ^45^. Identifying the loci underlying seed color can help breeders develop teff varieties that combine the resilience and nutritional benefits of brown seeds with the consumer preference and market value of white seededness.

Cultivated teff has striking variation in panicle architecture, ranging from very loose, open panicles resembling those of its wild progenitor *E. pilosa,* to semi-compact and fully compact forms similar to other cereals (Figure 4a). Compact panicles are generally more resistant to seed shattering and although they have improved harvestability and have generally lower yield. However, local farmer preferences and environmental conditions such as high humidity or traditional threshing practices can favor more open panicles, which dry faster and are easier to thresh manually. Consistent with breeding priorities, panicle architecture is strongly associated with seed color, with white-seeded varieties displaying significantly more compact panicles than brown-seeded varieties (Figure 4b).

**Figure 4.**
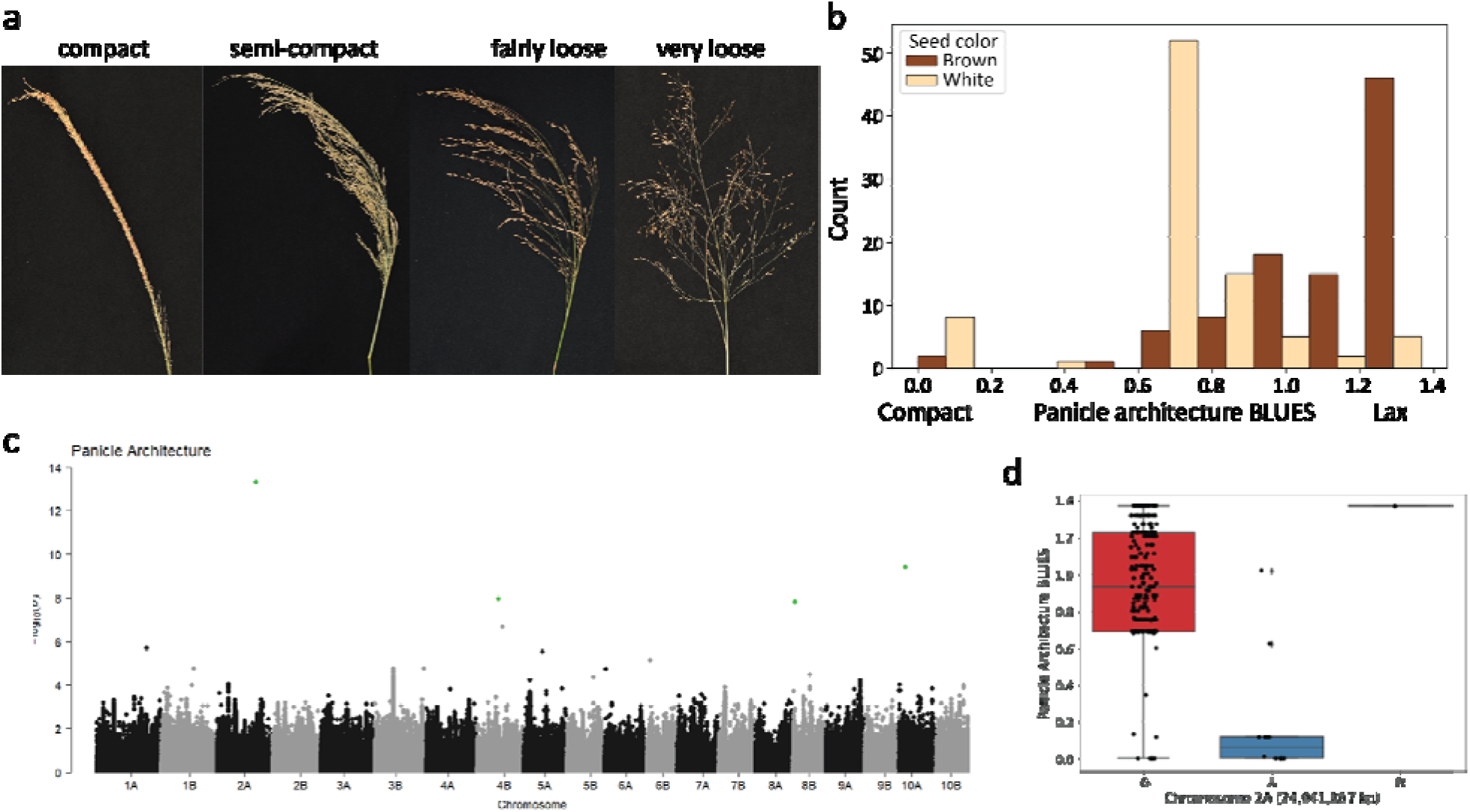
Genome-wide association analysis of panicle architecture traits in teff. (a) Representative teff panicles illustrating the range of panicle architecture, including compact, semi-compact, fairly loose, and very loose forms. (b) Relationship between panicle architecture (measured as BLUEs from field trials in 2021 and 2022) and seed color. BLUEs of 0 correspond to compact panicles, while values <1 represent increasingly open or lax panicles. (c) Manhattan plot showing results from genome-wide association analysis of panicle architecture BLUEs. Significant SNPs are highlighted in green. (d) Allele frequencies at the major locus on Chromosome 2A associated with panicle architecture variation.

We identified four loci significantly associated with panicle architecture in teff through GWAS of BLUEs from two years of field trials (Figure 4c). Given the linkage disequilibrium (LD) decay rates of 0.1 and 0.2 at 200 Kb and 68.5 Kb respectively (Figure 1c), we focused on candidate genes within a 200 Kb radius. The most significant peak was located on Chromosome 2A at 24,941,867 bp, 123 Kb from a previously reported GWAS hit associated with farmer preferences for panicle traits in the EtDP panel ^14^. This SNP falls within the second exon of Et_2A_018371, which encodes a 3-ketoacyl-CoA synthase-like protein with strong homology to the screw flag leaf1 (SFL1) gene in rice. Mutations in SFL1 cause a distinctive “screw” phenotype in rice, where both leaves and panicle branches exhibit helical twisting due to altered internode development ^46^. Interestingly, we observe a similar screw-like architecture in some compact teff panicles, suggesting possible functional conservation. In rice, *sfl1* mutants also show dn-type dwarfism, with uniformly shortened internodes.

A second significant GWAS hit was identified on Chromosome 4B at 13,698,024 bp, ∼180 Kb from Et_4B_037033, an ortholog of *dense and erect panicle* (*DEP1*) in rice and *erect panicle* (*EP*) in sorghum. These genes regulate inflorescence structure by shortening panicle internodes, increasing grain number per panicle, and promoting upright growth. DEP1 has been widely used in rice breeding to boost yield, while EP contributes to traits such as improved light interception, stronger stems, and enhanced photosynthetic efficiency from flowering through maturity ^47–50^. The proximity of these orthologs to teff GWAS peaks suggests that similar genetic pathways may underlie variation in panicle architecture across cereals. These candidate genes represent promising targets for marker-assisted selection to develop teff varieties with compact, high-yielding, and lodging-resistant panicles suited to both farmer preferences and mechanized harvesting.

Many teff varieties are grown as mixed genotypes with different seed colors. Within the TAP, 185 accessions produce brown or white seed consistently and these were used for GWA. Five loci were significantly associated with teff seed color, with two loci explaining 70% of the phenotypic diversity on Chromosome 4B (13,795,495 bp; 30.6%) and Chromosome 9B (1,536,847 bp; 39.6%) (Figure 5). Two alleles (C and G on Chromosome 9B and 4B, respectively) are found in 79% of the brown seeded varieties, and C and C alleles at these loci are in 91% of white varieties (Supplemental Figure 4). Interestingly, only the loci on Chromosome 4B overlaps with kmer based GWA hits for seed color identified in the Ethiopian diversity collection ^51^.

**Figure 5.**
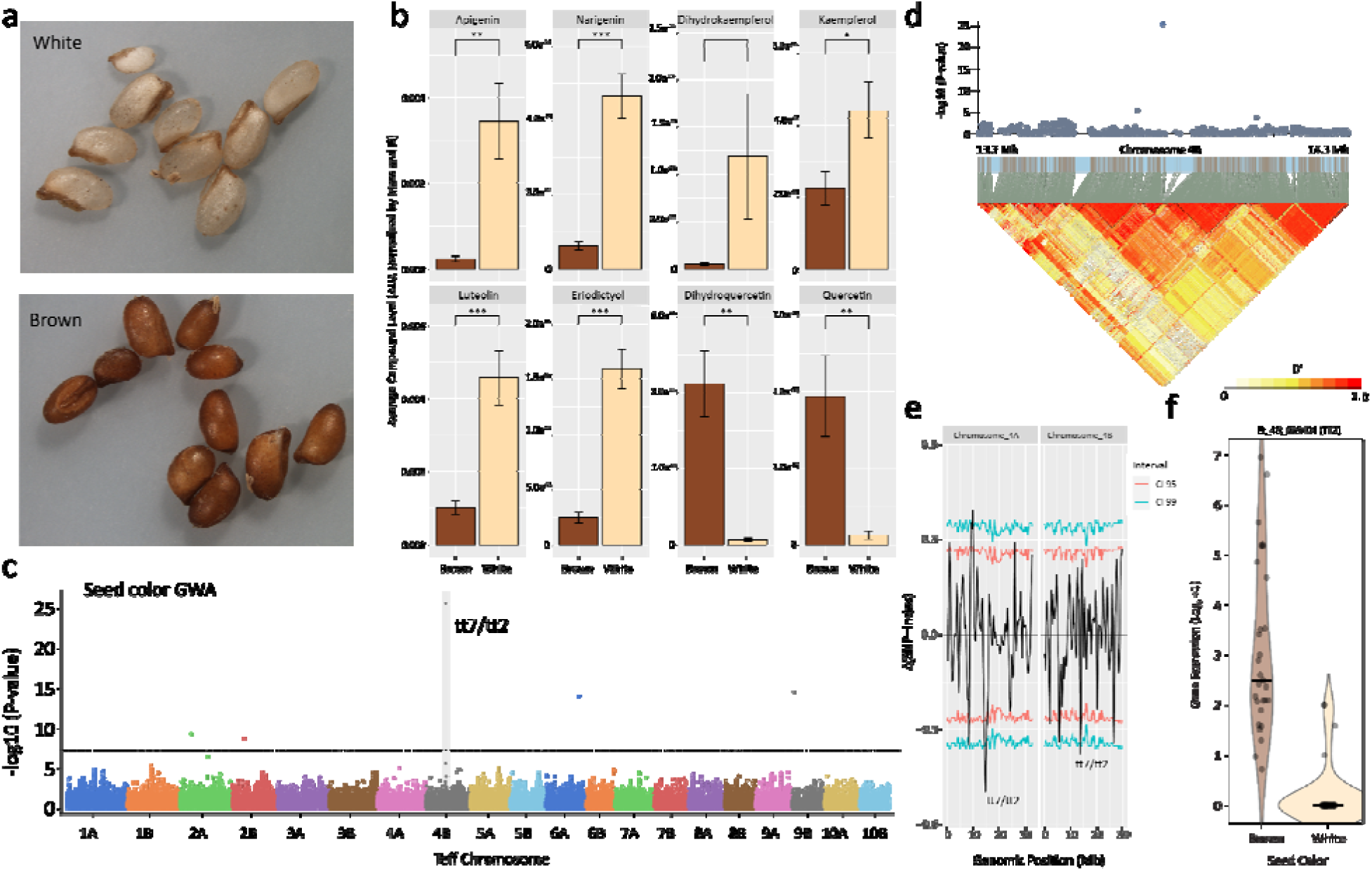
Identification of domestication genes related to seed color in teff. (a) Representative seeds from white and brown seeded teff varieties. (b) Log2-transformed concentrations of eight key flavonoids in brown and white teff seeds measured by UPLC-MS/MS. White seeds accumulate significantly higher levels of upstream flavonoids including apigenin, naringenin, dihydrokaempferol, kaempferol, luteolin, and eriodictyol, while brown seeds have significantly higher levels of downstream products dihydroquercetin and quercetin. Significance was determined using a two-sided t-test: p < 0.05 (*), p < 0.01 (**), p < 0.001 (***). (c) Manhattan plot from genome-wide association (GWA) analysis for seed color. The major loci on Chromosomes 4B containing orthologs to TT2 and TT7 is highlighted. (d) Local linkage disequilibrium (LD) heatmap showing strong LD surrounding the Chromosome 4B peak. (e) Bulk segregant analysis (BSA) of seed color using bulked F1 seeds from a single panicle. The plotted ΔSNP index (black line) represents allele frequency differences between bulked brown and white seeded individuals, with confidence intervals at 95% (red) and 99% (blue). The ΔSNP index is plotted for Chromosomes 4A and 4B and significant QTL are highlighted in the homeologous regions containing putative TT2/TT7 orthologs. (f) Violin plot of Log2 +1 transformed gene expression of the candidate TT2-like gene Et_4B_039404 in developing brown and white seeds. Each point represents a different replicate or timepoint from the developing seed datasets.

Et_4B_037025 is 79 Kb upstream of the loci on Chromosome 4B and is homologous to the Arabidopsis CYP75B1 gene, a flavonoid 3’-monooxygenase also known as transparent testa 7 (tt7). Arabidopsis *tt7* mutants exhibit a yellow seed coat due to the accumulation of kaempferol and loss of cyanidin-derived proanthocyanidins in the seed coat. This altered condensed tannin composition produces a yellow to pale brown seed coat compared to the dark brown of wild type^52,53^. A 1 bp deletion in the coding region of Et_4B_037025 is in linkage disequilibrium with the top GWA hit, and 87% of white accessions have the deletion and 87% of brown seeds have the reference genome ‘Dabbi’ allele. The homeolog on chromosome 4A of this gene has been lost in both the teff and *E. pilosa* genomes, and a single loss-of-function allele could be sufficient for a mutant phenotype.

Another candidate gene, Et_4B_039404, is an ortholog of *Arabidopsis TRANSPARENT TESTA2* (*TT2*), encoding a MYB transcription factor that regulates proanthocyanidin biosynthesis in seeds. It is located ∼200 Kb from the chromosome 4B peak SNP, within the extended LD block (Figure 5d). A long-terminal repeat (LTR) retrotransposon insertion in the first exon of Et_4B_039404 was previously identified in the white-seeded teff variety ‘Tsedey’, resulting in a truncated protein and likely loss of function ^51^. This structural variant could explain the white seed phenotype and may contribute to reduced proanthocyanidin accumulation through disruption of the TT2 regulatory pathway. To further support the role of TT2, we analyzed previously published RNA-seq data from developing seeds of brown and white seeded teff varieties ^13,54^. We found that expression of Et_4B_039404 is nearly undetectable in white seeds, but relatively high in brown seeds at several developmental stages (Figure 5f). In contrast, expression of the TT8 ortholog and many other regulatory and flavonoid biosynthetic genes is comparable between brown and white seeds (Supplemental Figure 5), consistent with the hypothesis that loss of TT2 function, rather than altered bHLH partner or pathway gene expression, underlies the white seed phenotype.

To independently validate these associations, we performed bulked segregant analysis using a teff accession which produces both white and brown seeds within a single panicle (PI 195933). Segregation was stable across two generations, and whole-genome resequencing of white and brown bulks identified 12 QTL regions containing 803 genes including 16 putatively involved in flavonoid, anthocyanin, or phenylpropanoid biosynthesis. QTL peaks on chromosomes 4A and 4B contain the homeologous TT2 orthologs identified via GWAS (Et_4A_032842 and Et_4B_039404; Figure 5e). The QTL in chromosome 4A has the highest significance and includes an InDel causing a frameshift in Et_4A_032842 that was also identified in the ‘Tsedey’ genome ^51^. Another QTL peak overlaps with the GWAS hit on chromosome 9B and includes Et_9B_066041 which encodes a putative anthocyanidin 5,3-O-glucosyltransferase. An A to G conversion on exon 1 caused a premature stop codon and disrupted the last 69 amino acids at C terminal.

Using a subset of white- and brown-seeded accessions, we conducted targeted reversed-phase ultra-performance liquid chromatography with tandem mass spectrometry (UPLC-MS/MS) to identify differentially accumulated flavonoid substrates and products between the two seed colors. Brown seeds had significantly higher levels of the flavonoids quercetin and dihydroquercetin (Figure 5b). In contrast, white seeds had significantly higher levels of apigenin, luteolin, naringenin, naringenin-7-*O*-glucoside, eriodictyol, and kaempferol (Figure 5b). No significant differences were observed for dihydrokaempferol, apigenin-7-O-glucoside, luteolin-7-O-glucoside, or eriodictyol-7-O-glucoside.

Previous in vitro functional characterization of flavonoid 3’-hydroxylases (F3’Hs) in other plant species suggests that apigenin, naringenin, dihydrokaempferol, and kaempferol can serve as substrates for the production of luteolin, eriodictyol, dihydroquercetin, and quercetin, respectively ^53,55,56^. If an F3’H knockout were responsible for white seed color, we would expect a reduction in F3’H products in these seeds. Given that brown teff seeds have significantly higher levels of the F3’H products quercetin and dihydroquercetin, and that dihydroquercetin is a known precursor for condensed tannin (proanthocyanidin) accumulation, this is a plausible explanation. However, the flavonoid profile of white seeds is also consistent with disruption of TT2, an R2R3-MYB transcription factor that regulates the late steps of proanthocyanidin biosynthesis in seeds. In Arabidopsis, tt2 mutants fail to activate DFR, ANS, and BAN, blocking the conversion of dihydroquercetin and related intermediates into catechins and proanthocyanidins. This block causes accumulation of upstream flavonoids such as naringenin, dihydrokaempferol, kaempferol, and various flavones and flavanones (apigenin, luteolin, eriodictyol), while downstream PA precursors and products are reduced. The high abundance of these upstream flavonoids in white teff seeds, together with reduced quercetin and dihydroquercetin, fits this metabolic pattern. Together, GWAS, BSA, expression, and metabolic data suggest a model in which independent mutations in TT2-like transcription factors contribute to natural variation in seed color in teff.

## Discussion and Conclusion

Here, we assembled and characterized the Teff Association Panel (TAP), a globally diverse set of teff landraces, cultivars, and breeding lines that captures much of the genetic diversity found across Ethiopia. Through resequencing and phenotyping, we confirmed the close evolutionary relationship between cultivated teff and its wild progenitor *Eragrostis pilosa*, and we leveraged this resource to dissect the genetic basis of key domestication and agronomic traits.

Our comparative genomic analyses reveal that *E. pilosa* and teff differ far less than expected for a domesticated wild species pair. The genomes are remarkably collinear, with minimal structural rearrangement, limited gene loss, and highly conserved gene content across both subgenomes. This level of conservation contrasts sharply with other cereal crops such as maize, sorghum, and rice, where domestication has been accompanied by widespread gene loss, structural variation, and dispensable gene content ^26–28^. This pattern is also unusual for an allopolyploid, where subgenomes typically experience homeologous exchange, gene fractionation, and subgenome dominance during domestication and improvement ^57–60^. In chloridoid grasses however, polyploidy is widespread ^61^, and the innate stability of their subgenomes, combined with small monoploid genome size and low transposable element content ^29,31,32,62^, may contribute to the genomic stability observed in teff. These findings suggest that teff was domesticated relatively recently and with limited genomic restructuring, possibly due to its high selfing rate, compact genome architecture, and a domestication process involving less intense selection pressure and weaker genetic bottlenecks than seen in other major cereals.

Domestication traits in teff, as in other cereals, encompass a range of morphological and quality attributes shaped by both agronomic demands and farmer preferences. Panicle architecture in teff spans a continuum from open, shattering-prone inflorescences resembling *E. pilosa* to compact, semi-erect panicles similar to other cereals. Compact panicles are often favored in formal breeding programs because they reduce shattering and improve mechanical harvestability. However, open or loose panicles remain the predominant type cultivated across most agroecological zones in Ethiopia, where they are generally regarded by farmers as higher yielding and well adapted to diverse environments. In contrast, compact panicle types are grown in more limited areas, sometimes valued for reduced shattering or easier harvest, but not as widely adopted. Our GWAS identified four loci associated with variation in panicle architecture, including candidate genes with known roles in inflorescence morphology such as orthologs to rice DEP1 and EP. Selection for seed color is similarly complex, as white seed is generally preferred by consumers for its culinary qualities and market value but brown seed varieties are often more resilient, higher yielding at elevation, and nutritionally enriched. Using GWAS, bulked segregant analysis, transcriptomics, and metabolomics, we identified multiple loci controlling seed color variation. Candidate genes include orthologs of TT2, which regulates proanthocyanidin biosynthesis in Arabidopsis; and the TT7 flavonoid 3’-hydroxylase, a key enzyme in anthocyanin biosynthesis. Variation in panicle architecture and seed color in teff is thus likely governed by a conserved set of orthologous genetic pathways shared across cereals.

Although we detected evidence of strong selective sweeps throughout the teff genome, none of these regions overlap with the major domestication loci we identified for seed color and panicle architecture. This suggests that key domestication traits in teff, such as panicle compactness and seed pigmentation, may not have undergone strong directional selection, or that multiple alleles and pathways contribute to their expression. In teff, where domestication appears to have been relatively diffuse and less intensive than in other cereals, selective sweeps may instead reflect partial fixation of domestication-related traits or the influence of local adaptation and farmer selection for regionally important traits. For example, loci associated with plant height, internode length, or drought resilience may have been shaped by environmental pressures or agronomic preferences, rather than classical bottlenecks associated with early domestication. This pattern is consistent with the high degree of genetic diversity retained in teff, the close genomic similarity to its wild progenitor *E. pilosa*, and the incomplete fixation of many agronomic traits.

Although teff retains several wild-type traits such as moderate shattering, variable lodging tolerance, and incomplete fixation of domestication alleles, our results demonstrate that significant improvements have already occurred since its divergence from *E. pilosa*. Notably, most genetic diversity remains structured among landraces rather than improved cultivars, suggesting that the full potential of teff’s gene pool has yet to be harnessed. Subpopulations 7 and 9, which cluster closely with *E. pilosa*, harbor high genetic diversity and could serve as reservoirs of useful alleles for traits such as stress resilience, yield, and nutrition. While the Ethiopian Institute of Agricultural Research has released dozens of improved teff varieties over the past two decades, these are concentrated in just two subpopulations, suggesting that broader representation of teff’s genetic diversity could enhance future breeding efforts. Many underutilized alleles in other subpopulations likely retain resilience traits while offering opportunities to improve agronomic performance. The TAP, coupled with genome-wide genotyping and a high-quality *E. pilosa* reference genome, provides a powerful foundation for modern teff improvement. These resources facilitate the identification of agronomically important loci and their deployment through molecular breeding and genome editing. By integrating genomic insights with local knowledge and farmer preferences teff breeding can be accelerated to meet growing demand for this climate-resilient and nutritionally valuable crop.

## Methods

### Plant Materials

The majority of the germplasm analyzed in this study was sourced from the USDA-ARS Germplasm Resources Information Network (GRIN; https://www.ars-grin.gov/). This included accessions of *Eragrostis heteromera* (4), *Eragrostis macilenta* (1), *Eragrostis mexicana* (2), *Eragrostis papposa* (4), *Eragrostis pilosa* (13), and *Eragrostis tef* (363). Our study surveyed all available teff accessions in the USDA GRIN available at the time. Three accessions of *Eragrostis pilosa* were acquired from the Royal Botanical Gardens, KEW germplasm database.

Some traditional or farmer-maintained accessions of teff consist of mixed lines and from these, we selected a single representative plant for sequencing. For whole genome resequencing, each plant was cultivated in a separate 4-inch pot under controlled conditions with a 14-hour light and 10-hour dark cycle at temperatures of 26°C during the day and 20°C at night in a greenhouse setting. A single leaf segment (approximately 50 mg) was harvested from one plant per accession and immediately stored at −80°C for subsequent DNA extraction.

### DNA Extraction and DNA-Seq Library Construction

For re-sequencing of teff accessions, DNA was extracted from leaves using the MagMax Plant DNA Isolation Kit (ThermoFisher # A32549). DNA concentration was measured using the Qubit HS DNA Kit (ThermoFisher # Q32854), and the quality of the DNA was verified by 0.8% agarose gel electrophoresis. Between 250-350 nanograms of DNA were used for DNA-seq library construction with the Kapa Hyper Plus DNA Kit (KapaBiosystems # KK8514), according to the manufacturer’s protocol with 6 PCR cycles. Normalized, multiplexed DNA-seq libraries were pooled and sequenced on the HiSeq4000 system in paired-end 150 nt mode at the Michigan State University Genomics Core.

### Read alignment and variant detection

Adapter sequences were trimmed from the paired-end reads using Trimmomatic v0.36. Reads shorter than 36 bp or with low-quality base pairs were removed. The trimmed reads were aligned to the *Eragrostis tef* genome assembly v3.1 ^13^ using Bowtie2 v2.2.3 using default parameters. The mean read alignment rates for all teff accessions is 98.8%. Teff is an allotetraploid, and the A and B subgenomes diverged an estimated ∼5 million years ago. There is little evidence of homeologous exchange, and the A and B subgenomes have an average nucleotide similarity of 93%, and we observed proper read alignment to the A and B subgenomes. The resultant SAM files were sorted by chromosome, read group information was added, and converted to BAM format using Picard Tools v2.18.27. SNP calling was performed with GATK v3.8, adhering to the GATK Best Practice protocols. HaplotypeCaller was utilized to genotype each accession, and all resulting VCF files were merged into a single file using CombineGVCFs, followed by joint genotyping with GenotypeGVCFs. The final VCF file underwent filtering to remove InDels and SNPs with a depth of coverage (DP) less than 10 and a quality by depth (QD) less than 30 using GATK’s SelectVariants function.

### Removing duplicated teff accessions

To detect closely related or nearly identical accessions, we performed an identity-by-descent analysis using PLINK v1.9. SNPs within linkage disequilibrium blocks were first pruned using PLINK with the option --indep-pairwise 50 10 0.5. Paired accessions with a PI_HAT value greater than 0.05 (indicating first to fourth degree relatives) were classified as cryptically related and grouped together. Within each cryptically related group, the accession with the highest number of sequencing reads was retained and the remaining accessions were removed from downstream analyses. All related accessions were identified exclusively within *Eragrostis tef* germplasm, with no related accessions found in other Eragrostis species or *Eragrostis pilosa*. In total, 135 accessions forming 36 groups were deemed related and 262 unique teff accessions were included for all downstream analysis. Our Teff Association Panel partially overlaps with the previously described Tef Diversity Panel of ∼300 accessions ^63^, but the two resources differ in sequencing strategy, filtering, and the final set of unique lines included.

### De novo assembly and annotation of the E. pilosa genome

Tissue for whole-genome sequencing was collected from a single immature *E. pilosa* plant from the seed collection at the Millennium Seed Bank (KEW0059857). Leaf tissue was flash-frozen in liquid nitrogen and stored at –80 °C prior to DNA extraction. High-molecular-weight genomic DNA was isolated by first extracting nuclei using the Circulomics Nuclei Isolation Kit, followed by DNA purification with the Circulomics Nanobind Plant Nuclei Big DNA Kit. PacBio HiFi libraries were prepared from the purified DNA and sequenced at the University of Georgia Genomics and Bioinformatics Core on a PacBio Sequel II platform. We generated 64.5 Gb of high-fidelity (HiFi) long-read sequencing data. Reads were assembled using hifiasm (v0.18)^24,64^ with default parameters and the number of haplotypes set to four (--n-hap 4). The resulting *E. pilosa* genome assembly consisted of 70 contigs totaling 560 Mb with an N50 of 14.7 Mb. Most chromosomes were assembled into 2–3 contigs. Full-length chloroplast and mitochondrial genomes were identified and retained, while partial or rearranged organellar sequences were removed.

Repetitive elements were annotated using the EDTA pipeline (v2.0.0) ^65^, which integrates HelitronScanner ^66^ for detection of DNA-based transposons and LTR_FINDER ^67^ and LTRharvest ^68^ for identification of long terminal repeat retrotransposons. Protein-coding genes were annotated using the MAKER-P pipeline (v2.31.10) ^69^, incorporating gene models from Oryza sativa ^70^, Arabidopsis thaliana ^71^, Oropetium thomaeum ^29,30^, and Eragrostis tef ^13^ as evidence. Ab initio gene prediction was performed using SNAP (2013 release) ^72^ and Augustus (v3.0.2) ^73^, with two rounds of iterative training to refine gene models. To remove gene models derived from transposable elements, we conducted a BLAST(v 2.10.0) ^74^ search against a curated non-redundant transposase database and filtered out matches. Assembly completeness was assessed using BUSCO (v2) ^75^ with the plant-specific embryophyte dataset. The resulting high-confidence gene models were used for all downstream analyses.

### Comparative genomics of the teff and E. pilosa genomes

Comparative genomic analyses between teff and *E. pilosa* were conducted using the MCScan toolkit (v1.1) ^76^ implemented in Python ([https://github.com/tanghaibao/jcvi/wiki/MCscan-(Python-version)]). Syntenic orthologs were identified with the chromosome scale teff ‘Dabbi’ genome ^13^ serving as the reference, anchor genome. Gene models from both species were aligned using LAST (v 914), and syntenic blocks were defined based on a minimum of five collinear gene pairs. Macrosynteny dot plots, depth histograms, and microsynteny visualizations were generated using the MCScan Python utilities. Conserved syntenic orthologs between teff and *E. pilosa* were extracted and used for downstream analyses of genome collinearity, gene retention, and polyploid subgenome comparisons.

To estimate evolutionary divergence, synonymous substitution rates (Ks) were calculated for homoeologous gene pairs within the *E. pilosa* subgenomes, and for syntenic orthologs between *E. pilosa* and teff. Ka and Ks values were computed using custom scripts available at [https://github.com/Aeyocca/ka_ks_pipe/]. Protein sequences for each gene pair were aligned using MUSCLE (v3.8.31) ^77^, and nucleotide alignments were generated with PAL2NAL (v14) ^78^. Ks values were then estimated using the codeml module from PAML (v4.9h) ^79^, with parameters specified in the control file provided in the GitHub repository.

### Nucleotide diversity estimation

We estimated nucleotide diversity (π) in teff and *E. pilosa* using the invariant sites of aligned reads to the teff reference genome using pixy (v1.2.7.beta) ^80^. For this analysis, we reran variant calling using mpileup in bcftools (v1.9.64) ^81^ using the sorted bam files as described above, with default parameters and on each chromosome separately. The resulting VCF file contained read depth for each individual for every base pair of the genome, providing a framework to estimate nucleotide diversity more accurately. Using pixy ^80^, we calculated nucleotide diversity (π) to estimate the genetic variation within each population, the fixation index (FST) to assess genetic differentiation among populations, and the average number of nucleotide substitutions per site between populations (Dxy) to understand the evolutionary distances ^80^. We calculated these metrics for all teff accessions vs. *E. pilosa* or for each teff subpopulation separately.

### Putative Sweep Identification

We identified genomic regions putatively under selection during domestication or improvement using three complementary approaches: cross-population composite likelihood (XP-CLR) ^82^, nucleotide diversity ratio (π; described above), and RAiSD (μ-statistic)^41^. XP-CLR was calculated using 8.1 million SNPs and the updated XP-CLR software ^82^ with a 50-kb sliding window and a 25-kb step size. The top 5% of XP-CLR scores were designated as candidate sweep regions. Nucleotide diversity (π) was calculated for both teff and *E. pilosa* as described above in 50-kb sliding windows, and regions with high π *E. pilosa* / π teff ratios were considered candidates for selective sweeps, reflecting reduced diversity in teff relative to its wild progenitor. In parallel, we assessed local deviations in the site frequency spectrum using RAiSD (μ-statistic)^41^ with default parameters and the same 8.1 million SNP file, also applied in 50-kb windows.

To identify high-confidence sweep regions, we retained genomic intervals that ranked in the top 5% of all the three methods (XP-CLR, π ratio, and RAiSD). Overlapping candidate regions were merged and used for downstream analysis, including annotation of genes within sweeps and assessment of functional enrichment.

### Locality analyses

Longitude and latitude for the TAP were collected from passport data on the NPGS GRIN-GLOBAL website (https://npgsweb.ars-grin.gov/gringlobal/search). Where locality was not provided, longitude and latitude were estimated based on information listed. EtDP longitude and latitude coordinates were collected from previous publication ^14^. Utilizing sf (https://r-spatial.github.io/sf/) a shapefile was created with longitude and latitude data and mapped to a shape file of Ethiopia from rnaturalearth. Precipitation and elevation were obtained from geodata worldclim and raster then visualized in R version 4.3.3.

### Bulk Segregant Analysis of Seed Color

Bulk segregant analysis (BSA) was conducted using teff accession PI195933, which produces both white and brown seeds. Seeds of each color were manually separated and grown in separate 3.5-inch pots under controlled conditions (14-hour light / 10-hour dark photoperiod, 25 °C day / 20 °C night). Leaf tissue from two-week-old seedlings derived from each seed color was pooled separately in 50 mL centrifuge tubes and flash-frozen in liquid nitrogen for DNA extraction. Three individual plants from each seed color group were maintained to maturity, and the seed color of their progeny was recorded. Genomic DNA was extracted from the pooled leaf tissue using the Omega Bio-tek E.Z.N.A. Plant DNA DS Kit (#D2411-01), following the manufacturer’s protocol. DNA-seq libraries were constructed and sequenced as described above. Libraries derived from white- and brown-seeded seedlings were designated “white bulk” and “brown bulk,” respectively.

Sequencing read trimming, alignment, and SNP calling were performed as described above. Variant call files (VCFs) were converted to tabular format using GATK’s VariantsToTable function to prepare input for the QTLseqr package in R ^83^. SNPs were filtered using the following parameters: minTotalDepth = 70, maxTotalDepth = 500, minSampleDepth = 70, depthDifference = 150, and minGQ = 99. Filtered SNPs were then analyzed using the G′ and Δ(SNP-index) methods implemented in QTLseqr, with a window size of 500,000 bp (5e5).

Twelve significant loci were identified in both analyses and selected for further investigation. Candidate genes within these 12 loci were examined for variants affecting pathways involved in anthocyanin, flavonoid, and phenylpropanoid biosynthesis and regulation. SNPs and InDels causing high-impact mutations such as frameshifts, gain or loss of stop codons, or changes in splice sites and showing an absolute Δ(SNP-index) > 0.5 between bulks were classified as strong candidate loci controlling seed color in PI195933.

### Multi-Year Field Evaluation of Morphological Traits in the TAP

Field phenotyping of the Teff Association Panel (TAP) was conducted in 2021 and 2022 at the Michigan State University Horticulture Teaching and Research Center in Holt, Michigan (42°67′43.4″N, 84°48′43.5″W). Each year, accessions were planted in triplicate in a randomized complete block design. Plots consisted of single 4.5-foot rows, with uniform spacing and management across the field.

To promote uniform growth and optimize yield, fertilizer (19-19-19 NPK) was applied pre-plant at a rate of approximately 100 lbs/acre. Weed control included broadcast application of Broclean for broadleaf weeds and targeted application of Roundup PowerMAX between rows for grass suppression. At maturity, panicle architecture was scored for each plot using a standardized 4-point scale: 1 = compact; 2 = semi-compact, 3 = fairly loose, 4 = very loose ^1^. After harvesting, seed from each plot was threshed, cleaned, and visually classified into three seed color categories: white, brown, or mixed.

### Genome wide association

Samples with consistent seed color of white and brown were selected for further analysis. Genome wide association was performed on 185 accessions for using Bayesian-information and LD iteratively nested keyway (BLINK) in GAPIT version 3 in R ^84,85^. The Best Linear Unbiased Estimators (BLUEs) for panicle architecture was estimated using lme4 where block and year were fit as random effects and accession was fit as a fixed effect, since plots were planted in a randomized complete block design. A kinship matrix and the first three PCs of the TAP PCA were included as covariates. Manhattan plots and QQ plots were constructed using QQman ^86^. Single nucleotide polymorphisms with a p value greater than Bonferroni corrected value were selected as significantly associated loci.

### Teff seed metabolite analysis

120mg ± 3mg of Eragrostis tef seeds were ground with a 5mm steel bead using a Fisherbrand™ Bead Mill 24 Homogenizer in bead mill tubes at speed 5, grind time 15 seconds, dwelling time 30 seconds, and 8 cycles. The samples were then centrifuged at 10,000 g for 60 seconds. 30mg ± 1mg of ground seed was then partitioned into new tubes with exact weights measured. 1000µL of 80% (v/v) HPLC-grade methanol and .1% (v/v) HPLC-grade formic acid was then added, and the samples were kept at 4°C for approximately 18 hours. The samples were then centrifuged at 13,000 g for 3 minutes at room temperature and 50µL of the supernatant was combined with 50µL of 50% (v/v) HPLC-grade methanol and .1% (v/v) HPLC-grade formic acid containing the internal standards picroside II, 8-prenylnaringenin, and phenylalanine d-8. 22 unique plots worth of seed representing 20 unique accessions with the seed from each plot divided into three technical reps. One plot (plot 2174) was divided into brown and white seeds with 17.1mg seed mass for the white seeds.

The samples were then analyzed using targeted liquid chromatography-tandem mass spectrometry (LC-MS/MS) with a Waters Acquity TQ-D UPLC-MS-MS equipped with a Waters Acquity Premier BEH C18 1.7µm 2.1 x 100mm column which was maintained at 40°C. The LC-MS/MS method used .1% (v/v) HPLC-grade formic acid as solvent A and 100% HPLC-grade acetonitrile as solvent B. The 10 min linear elution gradient at .300 mL/min consisted of 2% B held from 0 min to .5 min, 2%-50% B from .5 min to 6 min, 50%-99% B from 6 min to 7.1 min, held at 99% B from 7.1 min to 8 min, and held at 2% B from 8.01 min to 10 min. The samples were electrospray ionized in either positive or negative mode depending on individual compound ionization efficiencies in seven separate functions throughout each run. Blanks and quality control samples were included throughout the runs. The blanks contained 65% (v/v) HPLC-grade methanol and .1% (v/v) HPLC-grade formic acid with the internal standards. The quality control samples contained aliquots of 50µL from a pool which was created from 50µL of each seed extract. These aliquots were combined with 50µL of 50% (v/v) HPLC-grade methanol and .1% (v/v) HPLC-grade formic acid containing the internal standards. The resulting peaks were then identified by comparing to analytical standards. The values were determined by dividing the peak areas by the internal standard peak area within the run, then dividing that value by the seed mass used for that sample. Finally, these values were averaged between each of three technical replicates.

## Data Availability

Raw whole-genome resequencing reads for the Teff Association Panel and wild Eragrostis accessions have been deposited in the NCBI Sequence Read Archive (SRA) under BioProject PRJNA1321105. The *Eragrostis pilosa* reference genome assembly and corresponding gene annotations are available on CoGe.

## Acknowledgements

This work was funded, in part, by the Water and Life Interface Institute (NSF-DBI-2213983) to RV, the United States Department of Agriculture National Institute of Food and Agriculture (USDA-NIFA 2022-67013-36118) to RV and AT.

