## Supplemental Figures/Tables for "Origin, domestication, and diversity of the climate resilient Ethiopian cereal teff"

**Supplementary Tables and Figures**

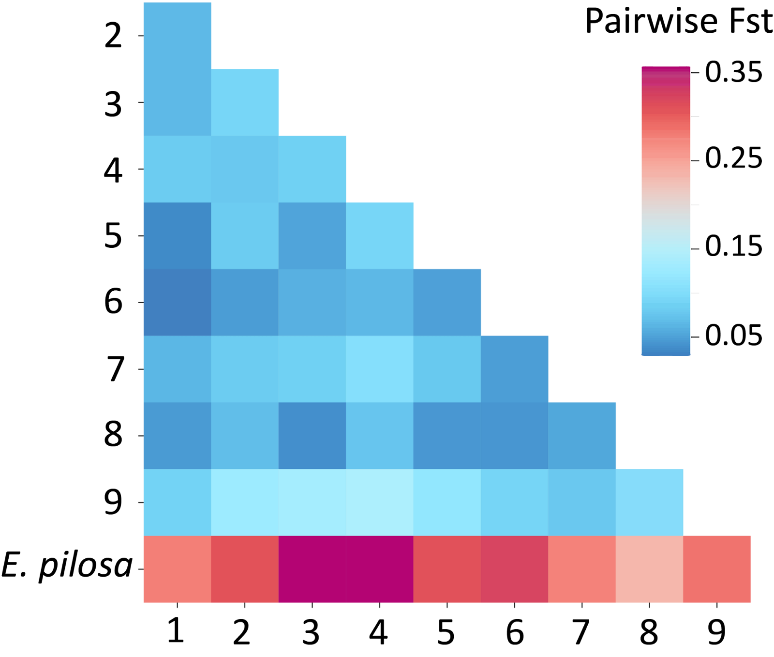

**Supplemental Figure 1. Pairwise population fixation (Fst) between accessions in each of the 9 teff subpopulations and *E. pilosa*.**

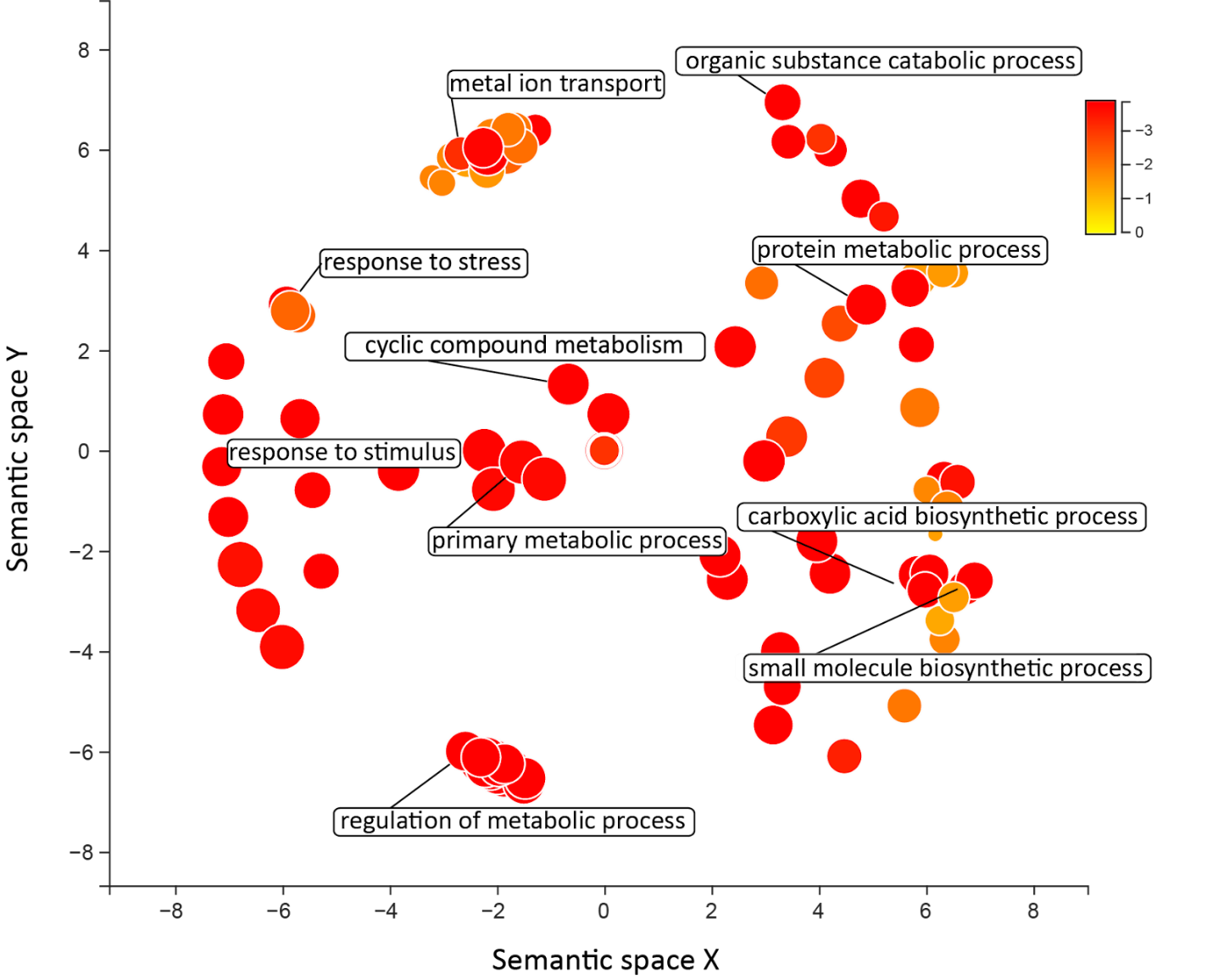

**Supplemental Figure 2. Enriched Gene Ontology (GO) terms of genes that are specific to teff.** GO terms are transformed using Multidimensional Scaling to reduce dimensionality and terms are grouped by semantic similarities. The color and size of the circles represent significance, and clustered processes of interest are highlighted.

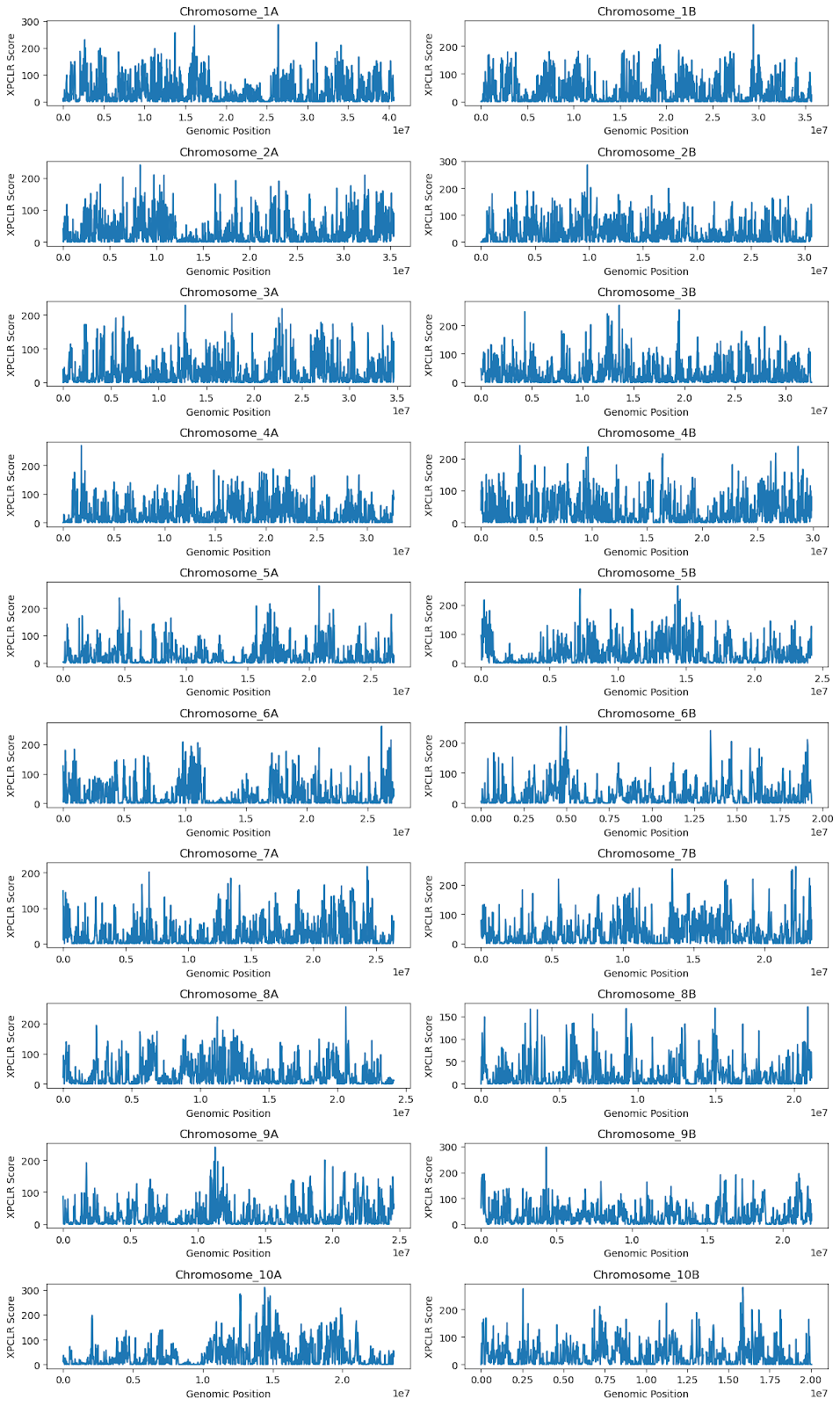

**Supplemental Figure 3. Genome-wide XP-CLR scan for selective sweeps in the teff genome.** XP-CLR scores were calculated in non-overlapping windows across each chromosome to detect regions under potential selection. Each panel represents a single chromosome, labeled from 1A to 10B, with the x-axis indicating genomic position (bp) and the y-axis showing the XP-CLR score. Peaks in the XP-CLR score correspond to candidate selective sweep regions. Homoeologous A and B subgenomes are displayed side by side to facilitate comparison of selective signals across subgenomic contexts.

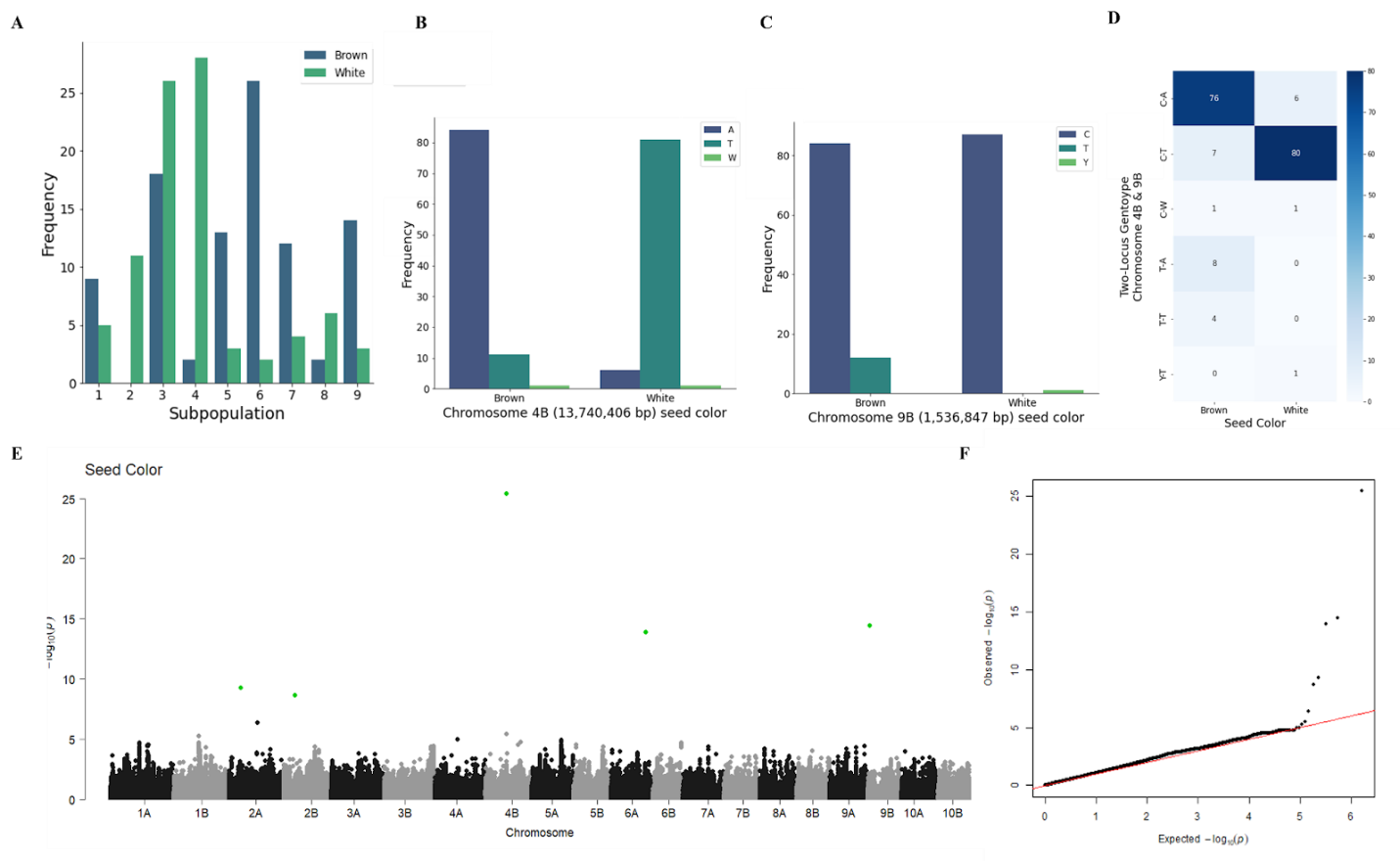

**Supplemental Figure 4. Seed color and genotypic association across the TAP. (**a) Seed color distribution across subpopulations. (b) Genotypic distribution at the peak Chromosome 4B SNP (position 13,740,406 bp) by seed color group. Most brown-seeded accessions are homozygous for the G allele, while white seeds are enriched for the C allele. (c) Genotypic distribution at the peak Chromosome 9B SNP (position 1,536,847 bp) by seed color, showing a similar association pattern. (d) Heatmap of genotypic combinations at the Chromosome 4B and 9B peak SNPs across accessions, stratified by seed color. The majority of brown seeded accessions carry the G/C combination, while most white seeded accessions are C/C.

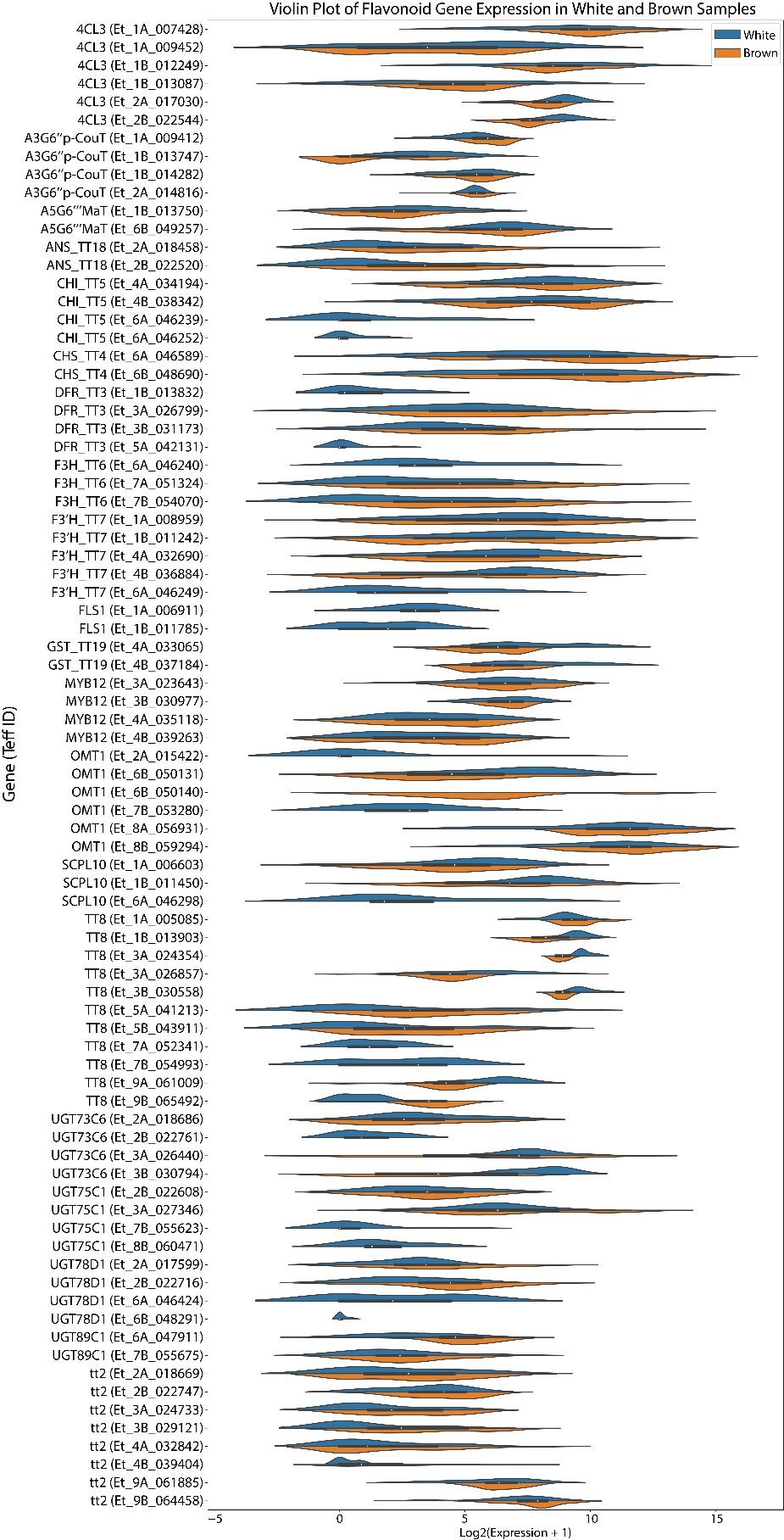

**Supplemental Figure 5. Expression of flavonoid biosynthesis pathway genes in brown and white teff.** Violin plot of Log2 +1 transformed gene expression from candidate flavonoid biosynthesis pathways genes collected from expression atlases of developing brown and white seeds.

**Supplemental Table 1: Sequencing read and alignment statistics of Eragrostis accession line used in this study. (see excel spreadsheet)**

**Supplemental Table 2: Summary of repetitive elements in the *E. pilosa* genome**

| **Class** | **Count** | **bp Masked** | **% masked** |
| --- | --- | --- | --- |
| **LTR** |  |  |  |
| Copia | 16940 | 20304144 | 3.62% |
| Gypsy | 41045 | 61122083 | 10.90% |
| Unknown | 18527 | 14441840 | 2.58% |
| **TIR** |  |  |  |
| CACTA | 25566 | 9603025 | 1.71% |
| Mutator | 42462 | 14832337 | 2.65% |
| PIF_Harbinger | 19972 | 6030512 | 1.08% |
| Tc1_Mariner | 20234 | 5244821 | 0.94% |
| hAT | 19677 | 7248824 | 1.29% |
| **nonTIR** |  |  |  |
| Helitron | 110770 | 47517078 | 8.47% |
| **total** | **315193** | **186,344,664** | **33.23%** |

**Supplemental Table 3. Summary of overlapping selective sweep regions in the teff genome.**

| **CHROM** | **window_pos_1** | **window_pos_2** | **xpclr** | **Pi Ratio** | **Mu_statistic** |
| --- | --- | --- | --- | --- | --- |
| Chromosome_10A | 12650001 | 12775000 | 283.59 | 24.84 | 5.72 |
| Chromosome_10A | 15375001 | 15425000 | 206.75 | 21.47 | 5.2 |
| Chromosome_10A | 15475001 | 15525000 | 157.41 | 24.46 | 6.24 |
| Chromosome_10A | 17050001 | 17100000 | 162.12 | 16.9 | 9.45 |
| Chromosome_10A | 18875001 | 18925000 | 156.13 | 25.54 | 5.67 |
| Chromosome_10A | 19325001 | 19375000 | 137.84 | 14.46 | 9.81 |
| Chromosome_10A | 19425001 | 19475000 | 172.1 | 26.46 | 7.92 |
| Chromosome_10B | 150001 | 200000 | 164.67 | 14.95 | 10.58 |
| Chromosome_10B | 16425001 | 16500000 | 198.88 | 19.78 | 5.53 |
| Chromosome_1A | 1075001 | 1125000 | 133.43 | 15.48 | 9.65 |
| Chromosome_1A | 1125001 | 1175000 | 137.25 | 23.46 | 10.54 |
| Chromosome_1A | 2575001 | 2750000 | 158.97 | 19.2 | 6.7 |
| Chromosome_1A | 5000001 | 5075000 | 126.02 | 15.84 | 5.48 |
| Chromosome_1A | 9650001 | 9850000 | 133.08 | 13.97 | 22.32 |
| Chromosome_1A | 12725001 | 12775000 | 156.57 | 14.38 | 5.23 |
| Chromosome_1A | 13725001 | 13800000 | 192.84 | 14.71 | 5.74 |
| Chromosome_1A | 15525001 | 15625000 | 125.51 | 18.03 | 5.79 |
| Chromosome_1A | 15675001 | 15725000 | 142.57 | 14.99 | 5.41 |
| Chromosome_1A | 15825001 | 15875000 | 125.06 | 22.56 | 7.21 |
| Chromosome_1A | 16425001 | 16500000 | 168.33 | 18.16 | 11.13 |
| Chromosome_1A | 33550001 | 33600000 | 133.62 | 18.58 | 6.87 |
| Chromosome_1A | 34050001 | 34175000 | 135.46 | 17.26 | 5.91 |
| Chromosome_1A | 38575001 | 38625000 | 141.52 | 17.87 | 8.56 |
| Chromosome_1B | 10100001 | 10150000 | 134.18 | 29.09 | 7.18 |
| Chromosome_1B | 10300001 | 10350000 | 153.28 | 14.7 | 8.34 |
| Chromosome_1B | 10400001 | 10450000 | 160.16 | 17.7 | 8.09 |
| Chromosome_1B | 18725001 | 18775000 | 154.31 | 14.89 | 6.3 |
| Chromosome_1B | 29900001 | 29950000 | 167.01 | 14.46 | 5.27 |
| Chromosome_2A | 2375001 | 2425000 | 134.97 | 14.89 | 5.78 |
| Chromosome_2A | 9775001 | 9825000 | 135.5 | 22.41 | 5.57 |
| Chromosome_2A | 10425001 | 10475000 | 126.19 | 26.11 | 6.77 |
| Chromosome_2A | 10800001 | 10850000 | 208.12 | 18.67 | 10.41 |
| Chromosome_2A | 30825001 | 30875000 | 136.57 | 15.6 | 5.25 |
| Chromosome_2A | 32225001 | 32275000 | 125.08 | 14.93 | 6.81 |
| Chromosome_2B | 775001 | 825000 | 130.15 | 18.78 | 7.53 |
| Chromosome_2B | 6825001 | 6875000 | 152.23 | 15.3 | 6.07 |
| Chromosome_2B | 8250001 | 8350000 | 159.66 | 14.74 | 5.34 |
| Chromosome_2B | 9700001 | 9775000 | 164.35 | 20.7 | 6.29 |
| Chromosome_2B | 10150001 | 10200000 | 201.94 | 16.04 | 7.61 |
| Chromosome_3A | 7125001 | 7175000 | 131.37 | 15.45 | 5.21 |
| Chromosome_3A | 22300001 | 22350000 | 160.59 | 14.13 | 7.82 |
| Chromosome_3A | 22650001 | 22750000 | 134 | 22.09 | 13.86 |
| Chromosome_3A | 26375001 | 26425000 | 132.21 | 14.58 | 11.02 |
| Chromosome_3A | 26575001 | 26625000 | 154.98 | 15.75 | 10.44 |
| Chromosome_3A | 27000001 | 27050000 | 177.5 | 23.1 | 5.17 |
| Chromosome_3A | 31900001 | 31975000 | 146.24 | 14.11 | 7.78 |
| Chromosome_3B | 2300001 | 2350000 | 157.36 | 16.03 | 7.38 |
| Chromosome_3B | 27925001 | 27975000 | 195.55 | 24.81 | 5.65 |
| Chromosome_4A | 1100001 | 1200000 | 152.74 | 19.27 | 5.99 |
| Chromosome_4A | 5025001 | 5075000 | 142.1 | 14.19 | 7.88 |
| Chromosome_4A | 19600001 | 19650000 | 177.75 | 28.74 | 6.08 |
| Chromosome_4A | 20050001 | 20100000 | 169.48 | 14.08 | 7.93 |
| Chromosome_4A | 20125001 | 20175000 | 148.76 | 19.14 | 7.93 |
| Chromosome_4A | 29200001 | 29250000 | 166.78 | 17 | 6.92 |
| Chromosome_4B | 7900001 | 7950000 | 127.39 | 15.19 | 5.55 |
| Chromosome_4B | 16375001 | 16475000 | 202.36 | 16.2 | 5.55 |
| Chromosome_5A | 16825001 | 16875000 | 215.7 | 16.1 | 5.18 |
| Chromosome_5B | 200001 | 325000 | 165.11 | 16.69 | 6.29 |
| Chromosome_5B | 400001 | 450000 | 176.08 | 15.19 | 6.35 |
| Chromosome_5B | 14750001 | 14800000 | 124.53 | 30.38 | 5.35 |
| Chromosome_6A | 4300001 | 4375000 | 136.74 | 17.22 | 5.89 |
| Chromosome_6A | 18975001 | 19025000 | 134.51 | 16.84 | 5.31 |
| Chromosome_6A | 26625001 | 26800000 | 178.67 | 31.14 | 5.34 |
| Chromosome_6A | 26900001 | 26950000 | 214.66 | 15.89 | 6.95 |
| Chromosome_6B | 19125001 | 19200000 | 210.15 | 27.98 | 6.75 |
| Chromosome_7A | 13175001 | 13225000 | 169.29 | 15.87 | 7.31 |
| Chromosome_7B | 10150001 | 10250000 | 167.37 | 14.97 | 7.6 |
| Chromosome_7B | 13475001 | 13575000 | 212.55 | 31.45 | 9.72 |
| Chromosome_7B | 14575001 | 14675000 | 168.05 | 14.28 | 5.87 |
| Chromosome_7B | 23175001 | 23225000 | 173.28 | 18.43 | 5.96 |
| Chromosome_8A | 6400001 | 6450000 | 130.56 | 15.32 | 10.58 |
| Chromosome_8A | 6475001 | 6525000 | 133.18 | 20.85 | 10.58 |
| Chromosome_8A | 11050001 | 11125000 | 151.25 | 26.25 | 8.09 |
| Chromosome_9A | 20850001 | 20925000 | 160.56 | 18.2 | 5.42 |
| Chromosome_9B | 1700001 | 1750000 | 137.38 | 15 | 5.24 |
| Chromosome_9B | 1850001 | 1900000 | 137.8 | 15.02 | 5.81 |
| Chromosome_9B | 11000001 | 11050000 | 162.91 | 13.94 | 5.36 |
| Chromosome_9B | 18050001 | 18100000 | 169.09 | 14.48 | 8.99 |

**Supplemental Table 4:** Percentage of brown and white seeded teff in each subpopulation included in GWA with 185 accessions.

| **Subpopulation** | **Percentage of Brown Seed** | **Percentage of White Seed** |
| --- | --- | --- |
| 1 | 64.3% | 35.7% |
| 2 | 0.00% | 100% |
| 3 | 40.9% | 59.1% |
| 4 | 6.67% | 93.3% |
| 5 | 81.3% | 18.8% |
| 6 | 92.9% | 7.10% |
| 7 | 75.0% | 25.0% |
| 8 | 25.0% | 75.0% |
| 9 | 82.4% | 17.6% |
